# Genome-wide association mapping of bruchid resistance loci in soybean

**DOI:** 10.1101/2023.09.22.559046

**Authors:** Clever Mukuze, Ulemu M. Msiska, Afang Badji, Tonny Obua, Sharon V. Kweyu, Selma N. Nghituwamhata, Evalyne C. Rono, Mcebisi Maphosa, Faizo Kasule, Phinehas Tukamuhabwa

## Abstract

Soybean is a globally important industrial, food, and cash crop. Despite its importance in present and future economies, its production is severely hampered by bruchids (*Callosobruchus chinensis*), a destructive storage insect pest, causing considerable yield losses. Therefore, the identification of genomic regions and candidate genes associated with bruchid resistance in soybean is crucial as it helps breeders develop new soybean varieties with improved resistance and quality. In this study, 6 multi-locus methods of the mrMLM model for genome-wide association study were used to dissect the genetic architecture of bruchid resistance using 5 traits: percentage adult bruchid emergence (PBE), percentage weight loss (PWL), growth index (GI), median development period (MDP), and Dobie susceptibility index (DSI) on 100 diverse soybean genotypes, genotyped with 14,469 single-nucleotide polymorphism (SNP) markers. Using the best linear unbiased predictors (BLUPs), 13 quantitative trait nucleotides (QTNs) were identified by the mrMLM model, 3 of which, rs16_14976250, rs1_22916615, and rs16_14975721, were associated with more than 1 bruchid resistance trait. As a result, the identified QTNs linked with resistance traits can be employed in marker-assisted breeding for the accurate and rapid screening of soybean genotypes for resistance to bruchids. Moreover, a gene search on the Phytozome soybean reference genome identified 27 potential candidate genes located within a window of 478.45 kb upstream and downstream of the most reliable QTNs. These candidate genes exhibit molecular and biological functionalities associated with various soybean resistance mechanisms and, therefore, could be incorporated into the farmers’ preferred soybean varieties that are susceptible to bruchids.

## 1. Introduction

Soybean (*Glycine max* (L.) Merr., 2n = 40) is an important food and cash crop worldwide with nitrogen-fixing ability and high oil and protein content (1). Despite its importance, the crop’s grain storage is affected by bruchids (*Callosobruchus chinensis*) (1), (2), (3). In most cases, the infestation starts in the field and spreads throughout the soybean value chain (3). Bruchids have high fecundity and the ability to re-infest, in addition to causing irreversible damage to soybeans (4); therefore, bruchid damage to soybean grain can lead to 100% yield losses (1), (3). To minimize yield losses to bruchids, farmers have been employing various management strategies including insecticides, botanically active plants, and cultural practices (5), (6). However, the majority of farmers tend to use insecticides for bruchid management (5). Despite their effectiveness, insecticides have several drawbacks such as risks to human and environmental health, high cost to resource-constrained farmers, and the possibility of the development of insecticide resistance (3), (7), (8). As a result, developing enhanced soybean cultivars with high resistance to *C. chinensis* is the most sustainable, cost-effective, strategically significant, and ecologically benign option.

For more effective development of soybean varieties that are resistant to *C. chinensis*, a thorough genetic dissection and understanding of the nature of this resistance is crucial. Recently, the study carried out by Msiska et al. (9) reported that soybean resistance to *C. chinensis* is due to the overproduction of secondary metabolites such as tannins and the high expression of enzymes such as peroxidases. A further genetic investigation of soybean resistance to *C. chinensis* revealed that the nature of gene action and mode of inheritance of bruchid resistance traits is quantitative and complex (10). Similarly, quantitative inheritance for resistance to bruchids has been reported in other legumes such as common bean (5) and cowpea (11). Breeding for such complex traits through conventional means is challenging due to environmental influences that slow down the selection progress. Hence, the use of molecular markers flanking the trait of interest improves the selection progress, although it necessitates a thorough genetic dissection and understanding of the association between the genotype and phenotypic variations (12). To the best of our knowledge, no study on association mapping and candidate gene discovery for soybean resistance to bruchids has been conducted in the soybean germplasm available at Makerere University Centre for Soybean Improvement and Development. This knowledge gap makes it difficult to implement marker-assisted selection (MAS) to combat the devastating impact of bruchids in Uganda and some parts of Sub-Saharan Africa that share the same genetic material. However, several studies have been carried out in other legumes aimed to determine quantitative trait loci (QTL) regions conferring bruchid resistance in cowpea using both bi-parental (13) and diverse (6), (12) populations and in common bean using bi-parental (14) and diverse (5) populations.

However, the use of bi-parental populations to identify QTLs has many drawbacks, including low resolution for QTL detection because only two parents contribute to the genetic information present in the mapping population, restricting its potential application in molecular plant breeding (6), (12), (15). To address this issue, genome-wide association studies (GWASs), a more robust contemporary mapping technique for identifying genomic regions and candidate genes linked with important traits in both self-pollinated and cross-pollinated crops, have been widely used (12). GWAS uses collections of lines with diverse genetic backgrounds and high historical recombination, resulting in more powerful QTL detection and better resolution (6), (12), (16). In soybean, GWAS has been used to identify genomic regions associated with resistance to many insect pests, including beet armyworm, potato leaf hopper, soybean looper, Mexican bean beetle, velvet caterpillar, and soybean aphid (17). Therefore, this study aims to identify quantitative trait nucleotides and candidate genes associated with resistance to *C. chinensis* in soybean using single-nucleotide polymorphism (SNP) molecular markers to uncover the genetic basis of bruchid resistance and facilitate MAS in soybean breeding.

## 2. Materials and Methods

### 2.1. Sources of Soybean Germplasm

The experiment consisted of 100 (80 resistant and moderately resistant and 20 susceptible) (S1 Table) soybean genotypes from different genetic backgrounds. These genotypes were obtained from SeedCo-Zimbabwe, Uganda, Japan, USA, IITA-Nigeria, and AVRDC-Taiwan.

### 2.2. Bruchid Rearing

Adult bruchid cultures were established in a laboratory at the Makerere University Agricultural Research Institute Kabanyolo (MUARIK) in 2018 (3). The bruchids used for culture establishment were originally obtained from the National Crop Resources Research Institute (NaCCRI) soybean storage in Namulonge, Uganda (3). Adult bruchids were allowed to oviposit on three susceptible commercially grown varieties (Maksoy 2N, Maksoy 3N, and Maksoy 4N) (3), with 1 and 5 kg of susceptible soybean seed placed in 1 L Kilner glass jars and 10 L plastic buckets, respectively, as described by Msiska et al. (3). Each jar and bucket were left at room temperature but covered with a muslin cloth to allow ventilation and the development of new adult bruchids while also preventing them from escaping. The bruchid populations were maintained by introducing them on a regular basis into new but susceptible soybean seeds, as described by Msiska et al. (3).

### 2.3. Bruchid Infestation and Phenotypic Data Collection

A sample of 100 soybean seeds was taken and weighed from each of the 498 genotypes to obtain the baseline for the 100 seed weight (3). To determine the initial seed weight, 50 randomly selected seeds from each soybean genotype were planted in various plastic Petri dishes and weighed. Following that, the soybean seeds in each Petri dish were artificially infested with 20 randomly selected adult bruchids from the bruchid colony using the no-choice test method (18) as described by Msiska et al. (3). The experiment was set up in a randomized complete block design, with insect infestation days serving as blocks, and it was replicated three times. The bruchids were kept in Petri dishes for 72 h to allow for mating and oviposition before being removed from the soybean samples after 10 days (3), (19). On day 11, the number of eggs laid on each of the 50 seeds was counted (3), and the number of emerged adult bruchids was counted and they were removed daily until no new insects emerged for 5 consecutive days (3), (20). The final weight of the seed samples in each Petri dish was recorded. Other bruchid resistance variables were estimated based on these data: Percentage bruchid emergence:

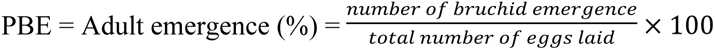

Percentage of weight loss:

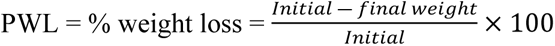

The growth index (GI), which is an indicator of genotype suitability for the development of insects, was calculated as follows:

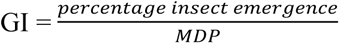

At the end of the experiment, the DSI (Dobie susceptibility index) was determined for each accession according to (21):

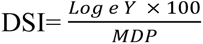

where *Y* = total number of emerging adults and MDP = median development period (days).

The median development period (MDP) was calculated as the number of days from the middle of oviposition (day 5) to the first progeny’s emergence (19) as described by Msiska et al. (2018). Kananji (19) suggested that DSI = 0, if no insects emerged over the test period. In this study, the modified susceptibility index ranging from 0 to 6 (3) was used to classify the soybean genotypes, where 0-1 =resistant, 2-3 = moderate resistant, 4-5 = susceptible, and 3-6 highly susceptible.

### 2.4. Phenotypic Data Analysis

To test soybean resistance to bruchids, the following parameters were used: number of eggs laid (NEL), percentage weight loss (PWL), Dobie susceptibility index (DSI), percentage adult bruchid emergence (PBE), median development period (MDP), and growth index (GI). Using the GenStat Statistical Package 18th Edition, each resistance parameter was subjected to a one-way analysis of variance (ANOVA). The Pearson correlation analysis was performed to determine the nature of relationships between bruchid resistance variables (3). All resistance parameters were analyzed using the GenStat Statistical Package 18th Edition following the linear model below:

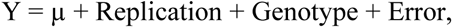

with Y being the phenotype of the target trait and μ its grand mean.

The broad-sense coefficient of genetic determination (H^2^) (i.e., equivalent to broad-sense heritability) was estimated as described by Piepho and Mohring (22) for all the soybean resistance traits using variance components obtained from a mixed model that considered the effects of all factors present in the model as random. The following formula was used:

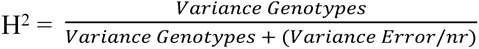

where *nr* is the number of replicates.

### 2.5. DNA Isolation, SNP Genotyping, LD Decay, and Multi-Locus GWAS Analysis

Seeds from 100 soybean genotypes were grown in a screen house at the Biosciences Eastern and Central Africa—International Livestock Research Institute (BecA-ILRI) Hub, Kenya, in 2019. Young, fresh leaf samples from each accession were collected twelve days after germination, and DNA was extracted using ZR Plant and Seed DNA Mini Prep^TM^ according to the manufacturer’s protocol with minor modifications (23). Prior to use, the DNA quality was determined on 0.8% (*w*/*v*) agarose gel in a 1 X Tris-acetate EDTA buffer and run at 80 V for 45 min, during which gel images were taken using a GelDoc-It^TM^ Imager (UVP), and the image was interpreted for DNA quality. Meanwhile, the DNA was quantified using a Thermo Scientific Nanodrop 2000C Spectrophotometer, Inqaba Biotech, South Africa and stored at 4 °C (23).

The DNA samples were sent to Australia, where the soybean genotypes were genotyped using the Illumina HiSeq 2500, Illumina Inc, California, USA and Diversity Arrays Technology Sequencing (DArTSeq^TM^), Australia. Subsequently, a genomic DNA library was constructed following an integrated DArT and genotyping-by-sequencing (GBS) methodology that involved complexity reduction of the genomic DNA, and repetitive sequences were eliminated using methylation-sensitive restrictive enzymes before sequencing on next-generation sequencing platforms (24).

The reads were aligned to the soybean reference genome Soybean_v7, which is publicly available at ftp://ftp.jgipsf.org/pub/JGI_data/phytozome/v7.0/Gmax (25), to identify single-nucleotide polymorphism (SNP) markers. Then, the HapMap genotypic data were determined for association analysis.

Prior to association analysis, the SNP data were filtered using a minor allele frequency (MAF) of 0.05 and a minimum count of 80% of the sample size using TASSEL v5.2.88 software (26). After filtering, a total of 14,469 SNPs were returned for subsequent analysis and imputed using the KNN imputation method implemented in TASSEL v5.2.88 (26), with all other parameters set to default. The software TASSEL v5.2.88 was used to calculate principal components (PCs) and the kinship matrix to infer the population structure and cryptic relatedness as described by Badji et al. (27). The kinship matrix was generated using the centered identity-by-state (Centered-IBS) function with other parameters set to default. The linkage disequilibrium (LD) decay, based on the squared Pearson correlation coefficient (r^2^) between pairs of SNPs, was calculated using TASSEL v5.2.88. An LD decay graph was generated by plotting the r^2^ between pairs of SNPs against their pairwise physical distance between pairs of SNPs as described in (28), (29) based on Remington et al. (30). The average pairwise distances at which LD decayed at r^2^ = 0.2 and 0.1 were also generated to visualize how LD decayed across the genome and allow the determination of the adequate LD blocks around markers.

For GWAS analysis, six multi-locus methods of the mrMLM package https://cran.r-project.org/web/packages/mrMLM/index.html, corrected for population structure and inequal kinship as described by Badji et al. (27), were applied to identify significant trait–marker associations using trait-phenotypic BLUPs and 14,469 imputed SNPs as described by Zhang et al. (31). GWAS analysis was performed on five bruchid resistance traits: percentage weight loss (PWL), Dobie susceptibility index (DSI), growth index (GI), percentage bruchid emergence (PBE), and median development period (MDP). Furthermore, both the kinship and the first 5 PCs were included in each of the six multi-locus methods (mrMLM, FASTmrMLM, pKWmEB, pLARmEB, FASTmrEMMA, and ISIS EM-BLASSO) in order to control false marker–trait association due to stratification effects and hence increase accuracy and power in quantitative trait nucleotide (QTN) detection (32). Additionally, the quantile–quantile (QQ) plots generated (S1 Fig) were used to visually assess the presence of spurious associations.

### 2.6. Annotation of Candidate Genes

The candidate genes underlying trait-associated SNPs involved in bruchid resistance functions were manually searched on the soybean reference genome hosted in Phytozome v13 https://phytozome next.jgi.doe.gov/geneatlas/jbrowse/index.html?data=genomes/Gmax_Wm82_a4_v1. Only QTNs that were detected by at least two multi-locus GWAS methods were considered for the candidate gene search. For this analysis, genes within a window of 478.45 kb upstream and downstream of the physical positions of the significant reliable QTNs, based on the genome-wide LD block size characterizing the mapping population, were searched on soybean reference genome and considered as candidate genes. Their functional annotations were closely examined from the Gmax_Wm82_a4_v1 soybean reference genome hosted on Phytozome v13, and their association with bruchid resistance was used as a prioritization criterion.

## 3. Results

### 3.1. Phenotypic Response of Soybean Accessions

Analysis of variance for soybean genotypes’ response to bruchid infestation showed significant differences (*p* < 0.05) for percentage weight loss (PWL), Dobie susceptibility index (DSI), percentage bruchid emergence (PBE), median development period (MDP), and growth index (GI) but not for number of eggs laid (NEL) (Table 1). Further, the broad-sense coefficient of genetic determination, which is equivalent to the broad-sense heritability, was relatively low to moderate for bruchid resistance traits: NEL (0.005), GI (0.10), PWL (0.31), DSI (0.22), PBE (0.21), and MDP (0.39) (Table 1).

**Table 1.** Analysis of variance with mean squares for soybean bruchid resistance.

| Source of Variation | d.f | PWL | NEL | PBE | DSI | GI | MDP |
| --- | --- | --- | --- | --- | --- | --- | --- |
| Genotypes | 497 | 108.31*** | 17.046<sup>ns</sup> | 13.08** | 4.2*** | 0.1011*** | 0.002737*** |
| Blocks | 2 | 4.47 | 2.34 | 2.38 | 0.999 | 0.4603 | 0.00153 |
| Residual | 998 | 74.97 | 17.38 | 10.39 | 3.26 | 0.3565 | 0.00166 |
| H<sup>2</sup> |  | 0.31 | 0.005 | 0.21 | 0.22 | 0.10 | 0.39 |

DSI = Dobie susceptibility index; PWL= percentage weight loss; NEL = number of eggs laid; PBE = percentage bruchid emergence; MDP = median development period (days), and GI = growth index. \* = $p < 0.05$ ; \*\* = $p < 0.01$ ; \*\*\* = $p < 0.001$ , and ns = not significant

The Pearson correlation analysis performed to determine the nature of relationships between bruchid resistant variables showed that number of eggs (0.78), PBE (0.87), and PWL were significantly positively correlated with DSI (*p* < 0.01), while weaker correlations with MDP were observed as described by Msiska et al. (2018). In addition, positive and significant (*p* < 00.1) correlations were observed among NEL, PBE, and PWL (S2 Table), suggesting that these bruchid resistance traits in soybean might be influenced by the same genetic basis.

### 3.2. Linkage Disequilibrium (LD) Decay

The genome-wide linkage disequilibrium (LD) computed for the 100 soybean lines using the 14,469 high-quality SNP markers is shown in Figure 1. The LD decay was relatively slow, with a distance of 478.45 kb at r^2^ of 0.2, decaying to an r^2^ of 0.1 at 1423.09 kb (Fig 1). This LD decay implies that the population is characterized by genomic LD blocks of 478.45, considering an LD threshold of r^2^ = 0.2.

**Fig 1.**
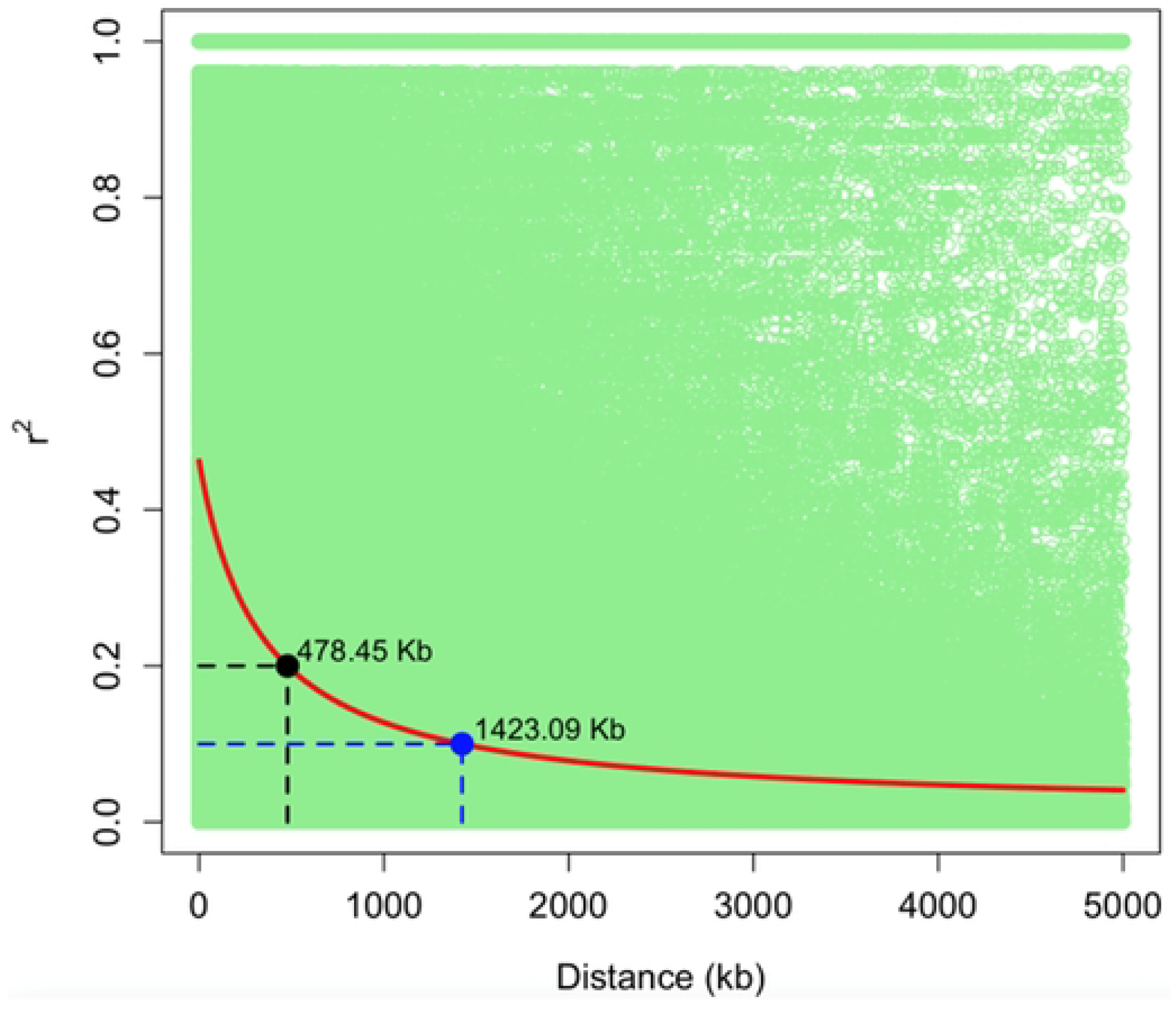
Linkage disequilibrium (LD) plot based on the 100 soybean lines and 14,469 SNP markers.

### 3.3. Association Mapping for Bruchid Resistance Traits

The results for the evaluation of the model fit for six multi-locus random-SNP-effect mixed linear models for GWAS (mrMLM) with the kinship and the PC matrices showed the best fit model to consider Q for all the bruchid resistance traits (PWL, GI, PBE, MDP, and DSI). Both the first 5 PCs and the kinship were included in the analysis to control false marker–trait association due to population stratification.

### 3.4. QTNs Associated with the Soybean Resistance Traits Identified by Multi-Locus GWAS

A total of 13 significant QTNs (LOD > 3) associated with soybean bruchid resistance traits were detected by 6 multi-locus GWAS methods across 7 chromosomes (Table 2 and Fig 2). Of these QTNs, rs7_14971060 on chromosome 7, rs18_14972895 on chromosome 18, and rs16_22914864 and rs16_14976250 on chromosome 16 had an allelic effect of 3.4995, 2.8772, 36906, and −0.0001 on PWL, respectively. The QTNs rs16_14976250 and rs16_14975721 on chromosome 16 and rs1_22916615 on chromosome 1 had an allelic effect of −29.6348, 17.6184, and 23.6492 on PBE, respectively. Furthermore, the QTNs rs15_14981757 on chromosome 15 and rs18_14978590 on chromosome 18 had an allelic effect of −1.5877 and −1.9444 on MDP, respectively. The QTNs rs13_14978774 and rs13_14973454 on chromosome 13; rs15_14981838 on chromosome 15; and rs16_22916222, rs16_22915136, and rs16_14976250 on chromosome 16 had an allelic effect of −1.1314, −3.00 × 10^−4^, 0.9582, 0.8024, 3.00 × 10^−4^, and −0.7096 on DSI. Three QTNs, rs16_14975721 and rs16_14976250 on chromosome 16 and rs1_22916615 on chromosome 1, associated with GI, had an allelic effect of 1.13 × 10^−5^, −0.566, and 1.5657, respectively (Table 2).

**Fig 2.**
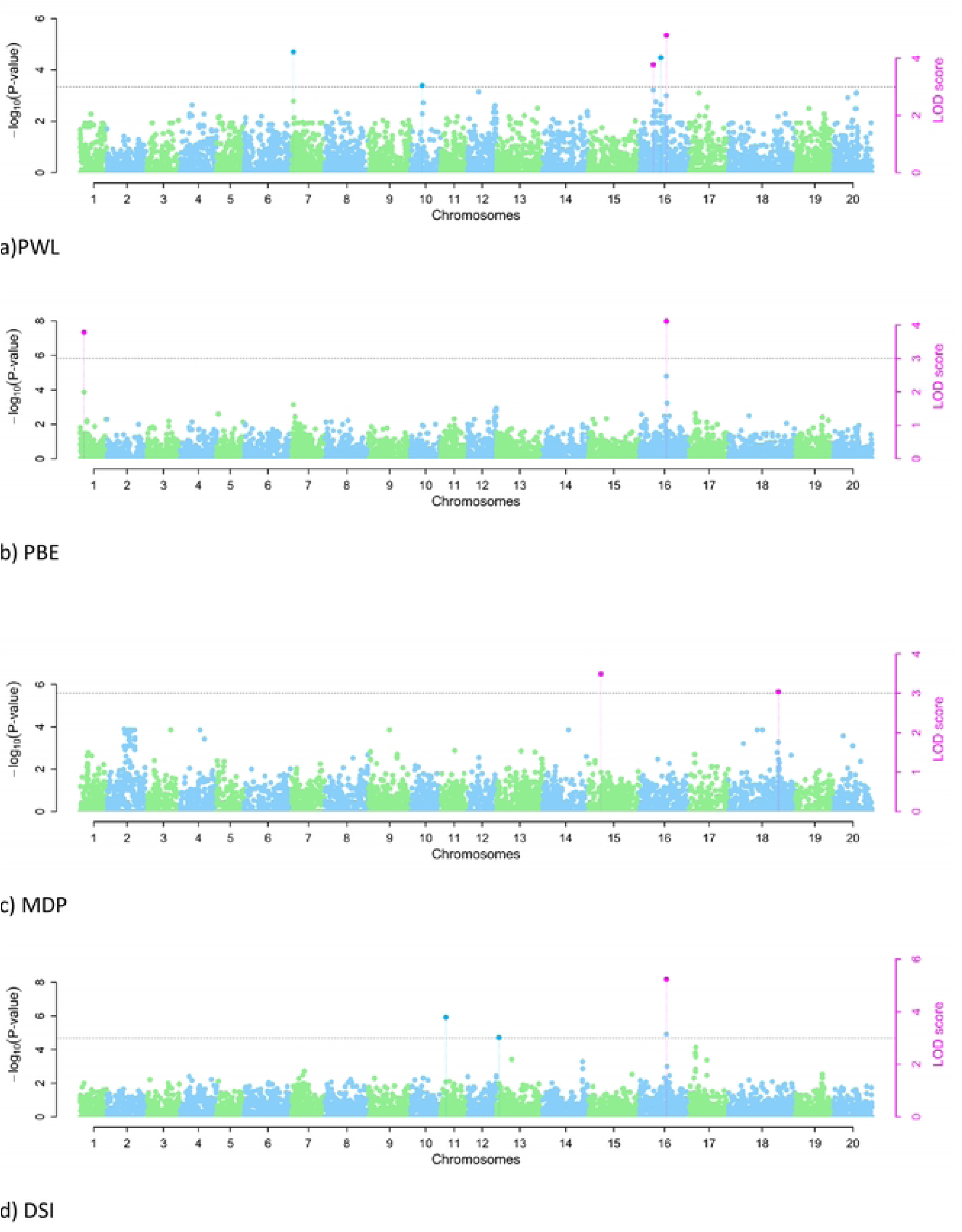
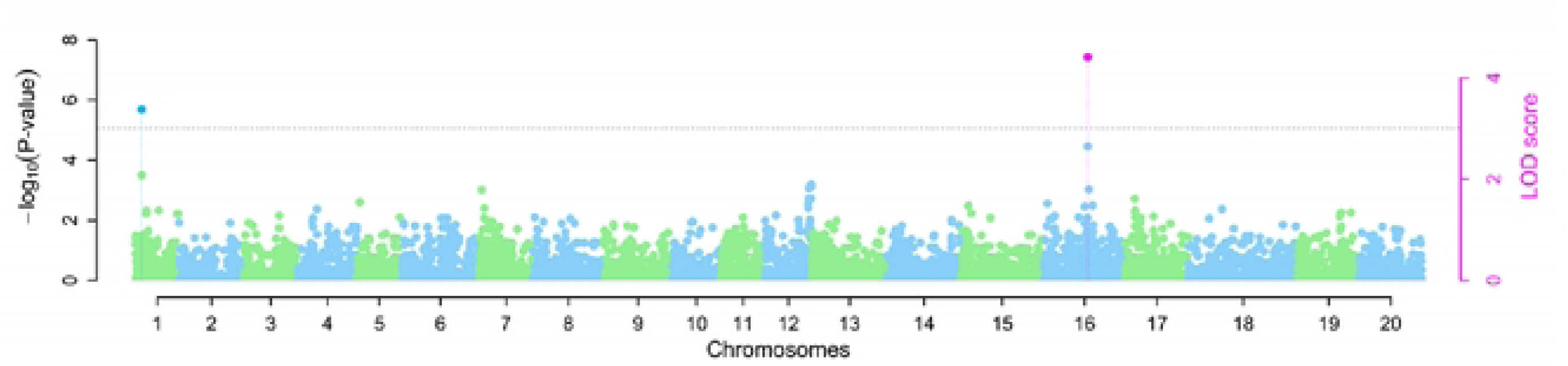
Manhattan plots for bruchid resistance traits in a subset of soybean mini-core collection. The horizontal dotted line indicates the genome-wide significance threshold (−log_10_ *p* ≥ 3.7). Association mapping of (A) PWL, (B) PBE, (C) MDP, (D) DSI, and (E) GI.

**Table 2.** Traits showing significant QTNs identified by one or more multi-locus GWAS methods.

| Trait | SNP | Chr | Marker Position (bp) | QTN Effect | LOD Score | R<sup>2</sup> (%) | MAF | Method |
| --- | --- | --- | --- | --- | --- | --- | --- | --- |
| PWL | rs7_14971060 | 7 | 2,312,825 | 3.4995 | 4.243 | 25.8378 | 0.4389 | 1 |
| PWL | rs18_14972895 | 18 | 54,931,863 | 2.8772 | 3.6637 | 11.6466 | 0.1889 | 1 |
| PWL | rs16_22914864 | 16 | 14,698,598 | 3.6906; 7.4002; 4.3707 | 4.1122; 4.0859 | 16.2244; 19.5078; 251,413 | 0.1739 | 3; 4; 5 |
| PWL | rs16_14976250 | 16 | 30,393,629 | −0.0001;<br>−2.3141 | 3.5023 | 3.22 × 10<sup>−8</sup>; 11.8927 | 0.4076 | 3; 5 |
| PBE | rs16_14976250 | 16 | 30,393,629 | −29.6348;<br>−20.4235;<br>−14.9328 | 4.2476; 7.3113; 4.2476 | 15.1816; 21.8395; 15.6575 | 0.4076 | 3; 4; 5 |
| PBE | rs16_14975721 | 16 | 27,267,774 | 17.6184 | 4.4654 | 10.0982 | 0.2283 | 5 |
| PBE | rs1_22916615 | 1 | 2,747,130 | 23.6492; 47.9168 | 3.6794 | 9.8751; 10.4641 | 0.0707 | 3; 4 |
| MDP | rs15_14981757 | 15 | 9,502,104 | −1.5877 | 3.4869 | 10.4855 | 0.3333 | 1 |
| MDP | rs18_14978590 | 18 | 55,103,397 | −1.9444 | 3.0341 | 8.7492 | 0.1413 | 3 |
| DSI | rs13_14978774 | 13 | 20,501,019 | −1.1314 | 3.3932 | 28.4305 | 0.1722 | 1 |
| DSI | rs15_14981838 | 15 | 3,067,634 | 0.9582 | 4.1483 | 19.702 | 0.3167 | 1 |
| **DSI** | rs16_22916222 | 16 | 29,157,714 | 0.8024 | 4.6506 | 14.5182 | 0.3333 | 1 |
| **DSI** | rs13_14973454 | 13 | 3,655,676 | $-3.00 \times 10^{-4}$ | 3.4329 | $2.08 \times 10^{-6}$ | 0.1576 | 2 |
| **DSI** | rs16_22915136 | 16 | 719,071 | $3.00 \times 10^{-4}$ | 3.091 | $2.70 \times 10^{-6}$ | 0.1902 | 2 |
| **DSI** | rs16_14976250 | 16 | 30,393,629 | -0.7096;<br>-1.4138;<br>-0.7096;<br>-0.7093 | 7.3247;<br>3.5839;<br>5.5482;<br>4.6236 | 21.4409;<br>20.9526;<br>19.4655;<br>21.4237 | 0.4076 | 2; 3; 4; 5 |
| **GI** | rs16_14975721 | 16 | 27,267,774 | $1.13 \times 10^{-5}$ | 4.38 | $4.47 \times 10^{-9}$ | 0.2283 | 2 |
| **GI** | rs1_22916615 | 1 | 2,747,130 | 1.5657 | 3.3427 | 9.0381 | 0.0707 | 4 |
| **GI** | rs16_14976250 | 16 | 30,393,629 | -0.566;<br>-0.4879 | 5.0186; 4.336 | 16.9309; 13.5202 | 0.4076 | 3; 5 |

Chr = chromosome; DSI = Dobie susceptibility index; PWL = percentage weight loss; PBE = percentage bruchid emergence; MDP = median development period (days), and GI = growth index. Methods: 1: mrMLM; 2: FastmrMLM; 3: ISIS EM-BLASSO; 4: FastmrEMMA; 5: pLARmEB; 6: pKMmEB.

The examination of QTN co-detection by multi-locus GWAS approaches revealed that four QTNs were simultaneously co-detected by more than two multi-locus methods (Table 2 and Fig 3). Among these QTNs, rs16_22914864 was associated with PWL and was simultaneously detected by ISIS EM-BLASSO, FastmrEMMA, and pLARmEB. The QTN rs16_14976250 was associated with four bruchid resistance traits, i.e., PWL, PBE, DSI, and GI, and simultaneously co-detected by FastmrMLM, ISIS EM-BLASSO, FastmrEMMA, and pLARmEB. The QTN rs16_14975721 was associated with PBE and GI and was co-detected by FastmrMLM and pLARmEB. The remaining QTN, rs1_22916615, was associated with PBE and GI, and was simultaneously detected by ISIS EM-BLASSO and FastmrEMMA (Table 2 and Fig 3). All the other QTNs were detected by only one multi-locus GWAS method; for instance, the QTNs rs7_14971060 (LOD score of 4.243) and rs18_14972895 (LOD score of 3.6637) were detected by mrMLM and explained 25.8378% and 11.6466% of the total phenotypic variation associated with soybean bruchid resistance trait PWL (Table 2 and Fig 3). The parameter PBE associated with the QTN rs16_14975721 (LOD score of 4.4654) was detected by pLARmEB and explained 10.0982% of the total phenotypic variation. The QTNs rs15_14981757 (LOD score of 3.4869) and rs18_14978590 (LOD score of 3.0341) accounted for 10.4855% and 8.7492% of the total phenotypic variation for MDP and were detected by mrMLM and ISIS EM-BLASSO, respectively. The QTNs rs13_14978774 (LOD score of 3.3932), rs15_14981838 (LOD score of 4.1483), and rs16_22916222 (LOD score of 4.6506), detected by mrMLM, and rs13_14973454 (LOD score of 3.4329) and rs16_22915136 (LOD score of 3.091), detected by FastmrMLM, were associated with DSI and accounted for 28.4305%, 19.702%, 14.5182 %, 2.08 × 10^−6^%, and 2.70 × 10^−6^% of the total variation, respectively (Table 2 and Fig 3). The GI was associated with the QTNs rs16_14975721 and rs1_22916615 detected with R^2^ values of 4.47 × 10^−9^ and 9.0381 by FastmrMLM and FastmrEMMA, respectively (Table 2). Interestingly, in this experiment, three QTNs were pleiotropic and were associated with more than one bruchid resistance trait (Table 2). The QTN rs16_14976250 on chromosome 16 was associated with the bruchid resistance traits: PWL, PBE, DSI, and G1, and it was detected by more than two multi-locus GWAS methods (Table 2). The QTN rs1_22916615 on chromosome 1 was associated with two resistance traits: PBE and GI, and it was detected by ISIS EM-BLASSO and FastmrEMMA (Table 2). The remaining QTN rs16_14975721 on chromosome 16 was associated with two bruchid resistance traits, PBE and GI, and detected by pLARmEB and FastmrMLM with R^2^ values of 10.0982 and 4.47 × 10^−9^ and LOD scores of 4.4654 and 4.38 for PBE and GI, respectively (Table 2).

**Fig 3.**
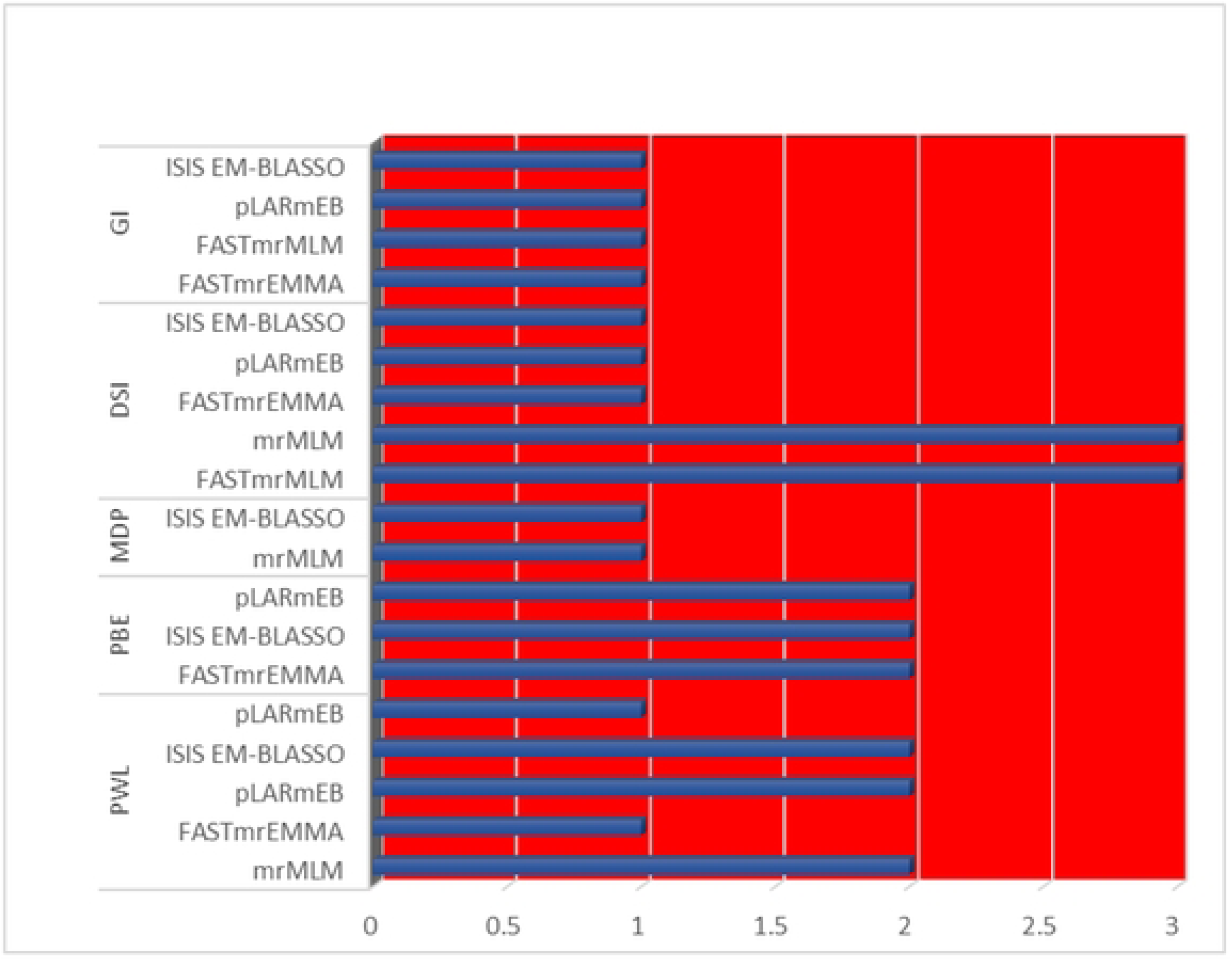
Number of significant QTNs detected by mrMLM model across bruchid resistance traits. DSI = Dobie susceptibility index; PWL = percentage weight loss; PBE = percentage bruchid emergence; MDP = median development period (days) and GI = growth index.

### 3.5. Candidate Gene Identification and Functional Annotation

Based on the LD decay, this study identified 27 putative candidate genes spanning the window 478.45 kb upstream and downstream of the most reliable QTNs (Table 3). The functionalities of these identified genes are known to be involved in plants’ responses to biotic and abiotic stress. Some of their notable functions in response to plant stress are through their involvement in MATE efflux family proteins, F-box family proteins, LIM domain-containing proteins, nucleic acid binding transcription factors, 2-oxoglutarate (2OG) and Fe (II)-dependent oxygenase superfamily proteins, MYB domain proteins, leucine-rich repeat (LRR) protein kinase family proteins, and GDSL-like lipase/acylhydrolase superfamily proteins (Table 3). The genes, *Gyma.16G140800*, *Glyma.01G025500*, *Glyma.01G026200*, *Glyma.01G026500*, *Glyma.01G027100*, *Glyma.01G027967,* and *Glyma.01G028700* found on chromosome 1 were associated with bruchid resistance traits, i.e., percentage adult bruchid emergence (PBE), and insect growth index (GI). Among these genes, *Glyma.01G026200* was very close to QTN rs_22916615, notably at 272 bp. In addition, *Glyma.16G091900*, *Glyma.16G092100*, *Glyma.16G092900,* and *Glyma.16G092600* found on chromosome 16, loci rs_22914864, were associated with percentage weight loss (PWL). Furthermore, on loci rs_14975721, the candidate genes *Glyma.16G121000*, *Glyma.16G119200*, *Gyma.16G124000*, *Gyma.16G124200*, *Gyma.16G120500*, *Gyma.16G122100*, *Gyma.16G121300*, *Gyma.16G123900,* and *Gyma.16G121700* were associated with the bruchid resistance traits PBE and GI. Moreover, on chromosome 16, *Gyma.16G140800*, *Gyma.16G141600*, *Gyma.16G142500*, *Gyma.16G142700*, *Gyma.16G143100*, *Gyma.16G144000,* and *Gyma.16G143102,* putative candidate genes involved in bruchid resistance traits, i.e., Dobie susceptibility index (DSI), PWL, PBE, and GI, were found in close proximity to the peak QTN rs_14976250 (Table 3).

**Table 3.**
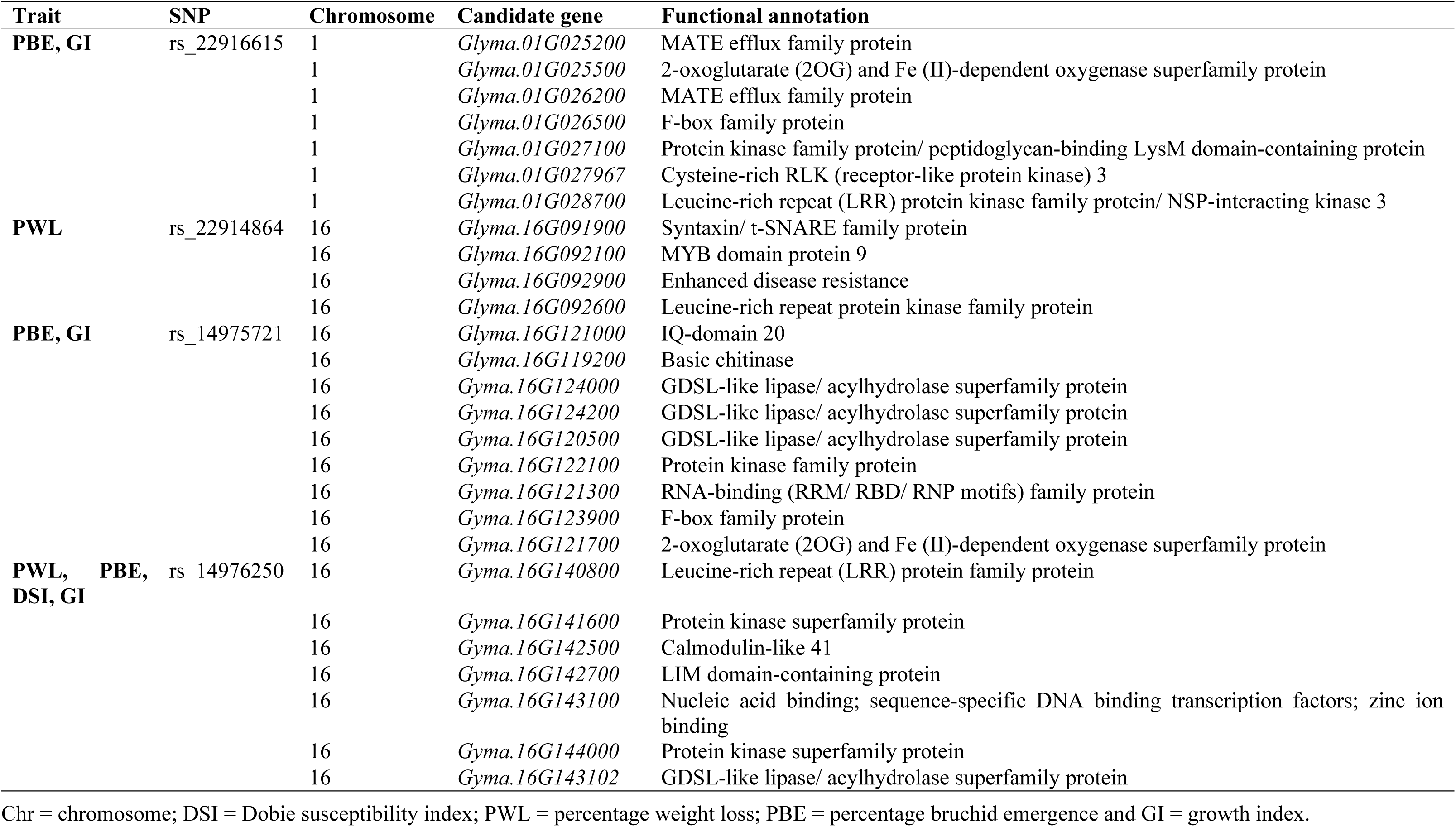
Annotation of identified candidate genes.

## 4. Discussion

The phenotypic evaluation for bruchid resistance traits showed highly significant differences among the genotypes evaluated. This implies the presence of genetic variation among the soybean genotypes and the possibility to conventionally identify and select genotypes for bruchid resistance in soybean; however, environmental influences could slow down the selection progress. The moderate heritability estimates obtained in this study were consistent with previous studies for bruchid resistance traits in cowpea (6) and common bean (19). These findings demonstrated the necessity of incorporating molecular techniques in the breeding of such complex traits to facilitate improvement.

Furthermore, the slow LD decay rate of 478.45 kb and the large LD block size observed in this study (Figure 1) could be attributed to the loss of genetic diversity caused by assortative mating during population improvement (33). Previous studies have reported slow LD decay in soybean: 220 kb for cyst nematode resistance (34), 544.01 kb for seed hardness (35), 138 kb for plant height and number of primary branches (36), and 200 kb for seed shape (37). Additionally, soybean is a strictly self-pollinated crop, which is expected to have a slower LD decay rate than outcrossing pollinated crops such as maize (33). However, this slow LD decay rate and large LD block size in soybean may lead to low resolution in association mapping and complicate candidate gene identification (33), (36).

The present study detected a total of 13 QTNs using multi-locus GWAS methods. The presence of significant QTNs associated with resistance to bruchids implies the predominance of additive gene action in conditioning soybean resistance to bruchids (10). Previous studies reported the predominance of additive gene action conferring resistance to bruchids in cowpea (11), (38) and common bean (19). This information could be useful for improving soybean resistance to bruchids through marker-assisted breeding. Furthermore, among the detected QTNs, three were associated with more than one trait and also linked to at least two candidate genes. This polygenic association between QTNs and numerous phenotypes and candidate genes confers a polygenic nature to inheritance in the regulation of bruchid resistance traits in soybean, which is of significant interest for improving farmer-preferred varieties that are susceptible to bruchids. The pleiotropic nature of genetic control for resistance to bruchids has been reported in cowpea (6), (39), (40).

The identification of candidate genes near reliable QTNs associated with traits of interest is considered a crucial post-GWAS analysis (27). In this study, 27 candidate genes associated with 4 bruchid resistance traits were identified within a window of 478.45 kb upstream and downstream of the 4 reliable QTNs on chromosomes 1 and 16. The functionalities of these identified genes are known to be involved in plants responses to biotic and abiotic stress. For instance, two upstream candidate genes, *Glyma.01G026500,* situated 20.9 kb of the peak QTN rs_22916615 on chromosome 1, and *Glyma.16G123900,* found at 363.4 kb of the peak QTN rs_14975721, are orthologous to *AT1G70590.1* and *AT4G35930.1,* respectively, in *Arabidopsis thaliana* and *Csa5M643280.1* in cucumbers, encoding for the F-box family protein, which plays a key role in jasmonate biosynthesis (41), (42), (43). Jasmonates play a crucial role in plants defense against insects, pathogens, and abiotic stresses (41), (44), (45), suggesting that these genes could also be involved in enhancing soybean resistance to bruchids.

In addition, two genes, *Glyma.01G025200* and *Glyma.01G026200,* were found in the vicinity of the peak QTN rs_22916615 (approximately 131.2 kb and 0.27 kb downstream, respectively) on chromosome 1. These genes encode the MATE efflux family protein transporters, which facilitate the intracellular transport of isoflavones in soybean (46). Isoflavones are major specialized secondary metabolites in soybean that play a crucial role in the plant’s response against pathogens and insect attacks (46), (47). Furthermore, we found four putative genes, *Glyma.01G027100*, *Glyma.16G122100*, *Glyma.16G141600,* and *Glyma.16G144000,* that encode protein kinase superfamily proteins (Table 3). In soybean, the protein kinase superfamily proteins play a key role in defense against insects such as aphids (48). Moreover, the other detected genes that could have contributed to bruchid resistance in soybean were *Glyma.01G02700*, *Glyma.16G092600,* and *Glyma.16G140800*. These genes are orthologous to *AT1G60800.1*, *AT5G48740.1,* and *AT3G50690.1* in *Arabidopsis thaliana*, encoding leucine-rich repeat (LLR) family proteins (Table 3). Leucine-rich repeat receptor-like kinase (LLR-RLK) is a family protein involved in plants’ responses to biotic and abiotic stress tolerance (49), (50). Also, *Glyma.16G142700* was detected on locus rs_14976250 just 34.9 kb downstream of the peak QTN. This gene encodes for ubiquitin-mediated regulated proteolysis, structural LIM domain-containing proteins, and the biosynthesis of brassinosteroid (BR), which plays a major role in seed coat development (51), (52). A hard seed coat would aid soybean defense against boring insect pests such as bruchids. Although seed hardening will be useful to avoid insect attack, it would not be useful for the soybean food industry; therefore, candidate gene prioritization is crucial. Interestingly, two candidate genes, *Glyma.16G121000* and *Glyma.16G142500*, detected on chromosome 16, encode calmodulin-like protein (IQ-domain 20 and 41, respectively) (53). Calmodulin-like proteins are known to play an essential role in plants defense against insects and pathogenic bacteria (53), (54), (55). Furthermore, some of the identified candidate genes are known to be involved in encoding for GDSL-like lipase/acylhydrolase superfamily proteins, nucleic acid binding transcription factors, basic chitinase, cysteine-rich RLK, MYB domain protein 9, and Syntaxin/t-SNARE family proteins and in the biosynthesis of phenolic acids and lignin, which all play a crucial role in the plant’s defense against biotic stress(56), (57), (58), (59), (60), (61), (62), (63), (64).

Given the plethora of candidate genes identified in this study, validation procedures are required to determine which genes are directly responsible for soybean resistance to bruchids. Moreso, QTNs associated with the candidate genes identified in this study need to be validated on different populations through gene knockout or fine mapping or gene expression studies. Furthermore, genetic information on resistance obtained through the validation process on different populations under local conditions would accelerate the advancement of marker-assisted breeding programs focusing on bruchid resistance in soybean varieties, particularly in Sub-Saharan Africa.

## 5. Conclusions

In the present study, 6 multi-locus methods of the mrMLM for GWAS identified 27 candidate genes that are closely associated with bruchid resistance in soybean. The identified candidate genes could be targeted for introgression in the development of new soybean varieties with improved bruchid resistance. Furthermore, the QTN rs_14976250, which was closely linked with major bruchid resistance traits, could be used for the fast and efficient breeding of soybean genotypes with a formidable bruchid resistance through targeted marker-assisted selection. The validation and implementation of the genetic information obtained in this study could be helpful in reducing postharvest loss due to bruchids in soybean.

## Supporting information

S1 Table. List and origin of 100 diverse soybean genotypes used in this study

S2 Table. Correlations among bruchid resistance traits

S3 Table. Summary of GWAS output for bruchid resistance traits

S1 Fig. QQ plots for GWAS for bruchid resistance traits.

## Acknowledgments

The authors would like to thank the Makerere University Centre for Soybean Breeding and Impovemnt for acquisition and provision of soybean germplasm from different regions. We are also grateful to the Biosciences Eastern and Central Africa—International Livestock Research Institute (BecA-ILRI) Hub for genotyping of the soybean population.

## Author Contributions

**Conceptualization**: Clever Mukuze, Ulemu M. Msiska, Tonny Obua and Phinehas Tukamuhabwa.

**Methodology**: Clever Mukuze, Ulemu M. Msiska, Tonny Obua

**Investigation**: Clever Mukuze and Ulemu M. Msiska

**Data curation**: Ulemu M. Msiska, Clever Mukuze and Tonny Obua

**Formal analysis**: Clever Mukuze, Afang Badji and Sharon V. Kweyu

**Funding acquisition**: Clever Mukuze

**Supervision**: Phinehas Tukamuhabwa and Mcebisi Maphosa

**Validation**: Clever Mukuze, Phinehas Tukamuhabwa, Ulemu M. Msiska, Tonny Obua, Afang Badji, Evalyne Chepkoech Rono, Selma N. Nghituwamhata, Faizo Kasule, Sharon V. Kweyu and Mcebisi Maphosa

**Writing - original draft**: Clever Mukuze, Mcebisi Maphosa and Faizo Kasule

**Writing - review & editing**: Phinehas Tukamuhabwa, Ulemu M. Msiska, Tonny Obua, Afang Badji, Evalyne C. Rono, Selma N. Nghituwamhata, Faizo Kasule, Sharon V. Kweyu and Mcebisi Maphosa

